# The Regime Shifts Database: A Framework for Analyzing Regime Shifts in Social-Ecological Systems

**DOI:** 10.1101/018473

**Authors:** Reinette (Oonsie) Biggs, Garry Peterson, Juan Carlos Rocha

## Abstract

This paper presents the Regime Shifts Database (RSDB), a new online, open-access database that uses a novel consistent framework to systematically analyze regime shifts based on their impacts, key drivers, underlying feedbacks, and management options. The database currently contains 27 generic types of regime shifts, and over 300 specific case studies of a variety of regime shifts. These regime shifts occur across diverse types of systems and are driven by many different types of processes. Besides impacting provisioning and regulating services, our work shows that regime shifts substantially impact cultural and aesthetic ecosystem services. We found that social-ecological feedbacks are difficult to characterize and more work is needed to develop new tools and approaches to better understand social-ecological regime shifts. We hope that the database will stimulate further research on regime shifts and make available information that can be used in management, planning and assessment.

## INTRODUCTION

Social-ecological systems can reorganize, unexpectedly shifting from being organized by one set of structures and processes to another. These regime shifts have been documented across an increasing range of systems ranging from the collapse of important fisheries to the salinization of agricultural soil (Gordon et al. 2008, Scheffer et al. 2001). Regime shifts impact the supply of essential ecosystem services on which human societies depend, such as crop production, flood regulation, and opportunities for recreation. These changes can have major impacts on human economies, security and health (MA 2005, Rocha et al. 2015). Better understanding the potential risks and consequences of regime shifts has been identified as an urgent priority (Carpenter et al. 2009, Reid et al. 2010), particularly in the context of processes such as Future Earth, the Intergovernmental Panel on Climate Change (IPCC), and the Intergovernmental Platform on Biodiversity and Ecosystem Services (IPBES).

This paper introduces a new database, the Regime Shifts Database (RSDB), which is aims to support such initiatives by providing policy-relevant information on regime shifts, and using a consistent framework to systematically analyze their impacts, key drivers, underlying feedbacks, and management options. Most regime shift research has focused on the analysis of individual regime shifts rather than comparison across multiple regime shifts. Although there have been several system-specific reviews addressing certain domains (Gordon et al. 2008, Lenton et al. 2008, Mård Karlsson et al. 2011), no comparative cross-system synthesis of regime shifts across multiple system types currently exists. This lack of comparative studies is partly due to a lack of agreement on practical, operational criteria for identifying regime shifts across different system types and different disciplines.

Researchers working on regime shifts are united by their focus on understanding large, abrupt, persistent systemic change. Although such regime shifts occur rarely, they are often highly relevant to policy due to the magnitude of their impacts, their persistence and their unexpected nature. The key debate in regime shift research is whether a given change is indeed a large, persistent, systemic reorganization of a system. Large, abrupt changes can arise from a sudden change in a system driver, or from a non-linear relationship between a key driver and response variable, and do not necessarily indicate a reorganization of internal feedback processes within a system that cause a system to persist in another state (Andersen et al. 2009). In fields such as oceanography large empirically observed step changes in ecosystem or social-ecological responses are regarded as regime shifts (Anderson and Piatt 1999), while in fields such as ecosystem ecology and earth system science much more emphasis is placed on whether system feedback processes are moving the system towards alternative attractors (Biggs et al. 2012, Scheffer et al. 2001). In empirical cases with limited understanding and noisy data it can be difficult to distinguish different types abrupt change from regime shifts.

However, from the point of view of the people living within a social-ecological system, the impacts of a large, abrupt, persistent systemic change are often identical regardless of whether it represents an true alternative theoretical equilibrium or a transient state that persists over a societally long term timescale (e.g. years to decades). It usually takes decades to clearly establish whether processes underlying an observed change are capable of generating alternative theoretical attractors (e.g., Schindler 2006), and due to internal and external system dynamics the existence of possible theoretical attractors may also vary over time scales comparable to the dynamics of the system (Biggs et al. 2009). In many situations it is useful to simply know that a system can experience substantial, surprising, persistent change even if the exact nature of that change is unknown.

The Regime Shifts Database (RSDB) adopts a pragmatic human-centered approach to regime shifts. We focus on identifying potential regime shifts, and documenting uncertainty surrounding their dynamics, rather than attempting to resolve theoretical issues. The RSDB draws strongly on a systems-based understanding of the dynamics underlying regime shifts (Box 1). It focuses on examples of large, persistent systemic changes in SES that impact ecosystem services, for which there are established or at least proposed changes in internal feedbacks that sustain the change and make it difficult to reverse. Our goal is to translate the concept of regime shifts into a pragmatic approach that can usefully inform regime shift policy, planning, assessment, and management in the face of incomplete evidence and information in a rapidly changing world. We hope that the RSDB will provide a platform for future meta-analyses of regime shifts and facilitate knowledge transfer between more and less well-studied regions of the world. The RSDB is inspired by the Resilience Alliance’s Threshold Database (Walker & Meyer 2004) as well as large cross-case syntheses on the management of communal forests (Moran and Ostrom 2005) and drivers of tropical deforestation (Geist and Lambin 2002), and hopes to advance understanding of regime shifts in SES in a similar way.

#### Box 1: Understanding regime shifts: A Systems View

**Figure B1.**
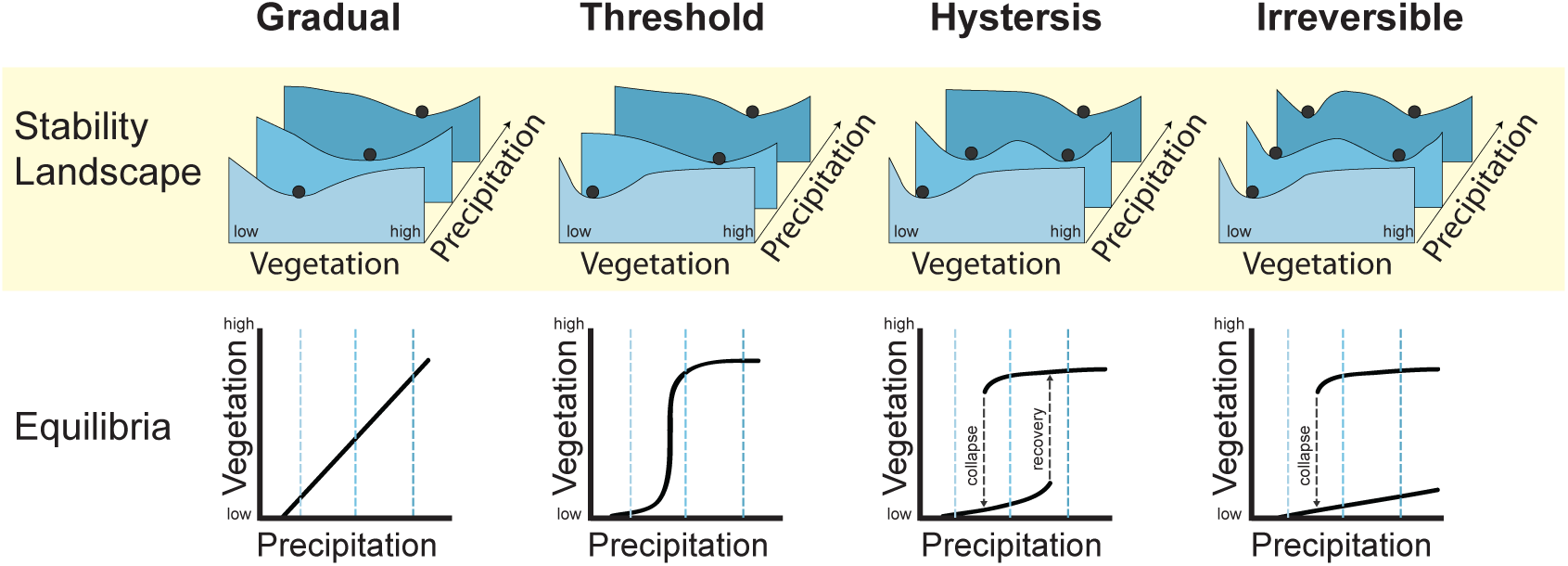
Stability landscapes for different types of ecosystem change. Here as an example the state or response variable of the system is precipitation while the slow variable is precipitation. The system can respond in a gradual or semi-linear fashion, abrupt jumps following thresholds, showing difficulties to reverse when hysteresis is at place, or irreversible change.

Ecology has often used a systems perspective to analyze the organization of nature (ONeill, 2001). In late 60’s there was a fundamental paradigm shift in ecology. Ecosystems were thought to tend towards punctuated equilibrium, but it was demonstrated that ecological systems tend to self-organize into configurations that allow variation and fluctuations on species composition, and yet maintain resilience and identity (Holling 1973, May 1977). Ecosystem stability is characterized by multiple domains or attraction, also know as regimes. One of the main ways of analyzing such systems is to identify the feedback loops that regulate the dynamics of a system. Systems can typically be viewed as consisting of sets of linked elements by feedback loops. As a system changes, different feedback loops tends to become dominant, so that the system becomes structured and functions in a particular way – forming a particular regime. Dominant feedbacks tend to be self-reinforcing, creating conditions that enhance their persistence, and making regimes “sticky” once they form.

A regime shift occurs when a switch in the dominant feedbacks occurs, and is often associated with rapid non-linear change as the system reorganizes into a different structure and function. Such a switch can occur when a large shock (e.g. hurricane) overwhelms the dominant system feedbacks. More commonly, a gradual change (e.g. habitat loss) slowly erodes the strength of the dominant feedbacks until a threshold is reached at which a different set of feedbacks suddenly becomes dominant and the system rapidly becomes reorganized into a new regime. Drivers of regime shifts are usually understood as external variables to the system dynamics that directly or indirectly influence its dynamics (Nelson et al. 2006). Regimes are understood as the region of the parameter space (e.g. vegetation cover vs precipitation) where the dynamics of the system tend to fluctuate. Regimes, basins of attraction or alternative stable states are all concepts that denote this particular region.

If a regime shift occurs the system reorganizes into a different configuration, i.e. a different set of dominant components and feedbacks amongst these, resulting in a different structure and function of the system (Biggs et al. 2012). The slow erosion of feedbacks usually goes unnoticed until the actual regime shift occurs – hence the shift often comes as a surprise. Furthermore, because the new dominant feedbacks are also self-reinforcing, the shift may be costly or impossible to reverse (Scheffer 2009, Scheffer *et al*. 2001). However, this is not always the case, by implementing managerial options on key elements of the systems, its behavior can be restored towards a societal desired regime. Such points of intervention are also known as leverage points (Meadows 2008).

This paper introduces the RSDB in four stages. We first describe the database and the criteria we use for selecting examples for inclusion. Second we describe the framework for documenting regime shifts that we developed for analyzing examples of regime shifts. Third, we present a synthesis of the examples included in the RSDB to date, highlighting interesting emerging patterns and possible avenues for further research. We conclude with a discussion of what we have learnt from the development of the framework to date.

## THE REGIME SHIFTS DATABASE (RSDB)

The RSDB systematically compiles examples of regime shifts in social-ecological systems that have large consequences for ecosystem services and human well-being in order to provide novel empirical and theoretical syntheses that can inform and support the emerging resilience and sustainability science fields, and initiatives such as Future Earth. The target audiences for the RSDB are researchers, lecturers and practitioners concerned with issues of environmental change.

The database is freely available online at www.regimeshifts.org, and aims to provide a platform that can serve as an entry point to identify policy-relevant regime shifts for purposes of research, teaching and environmental management. The database can, amongst others, be searched for regime shifts influenced by a particular driver, occurring in a particular ecosystem or land use type, or that have specific ecosystem service or human well-being impacts. Each entry also includes an explanation of the underlying drivers and dynamics that lead to the shift. A variety of open source material such as images and toy models are available for teaching purposes. This information may be directly used by lecturers or practitioners engaged in ecosystem or resilience assessments (Ash et al. 2010, RA 2007), or provide a starting point for more detailed scientific analyses, for instance of drivers of regime shifts (Rocha et al. 2015).

The database contains three “levels” of regime shifts. The first is different “generic types” of regime shifts, such as lake eutrophication or bush encroachment. These are general syntheses of regime shifts that have been observed in many localities around the world. The second and third levels are detailed and basic case studies of regime shifts in specific places. The detailed case studies provide substantial information and analysis of a specific regime shift, for example in the Baltic Sea. The basic case studies simply provide a short description of a case and links to reference. The examples included in the RSDB are derived from the literature, particularly synthetic reviews of regime shifts in particular systems (e.g., Gordon et al. 2008, Lenton et al. 2008, Nyström et al. 2012), and also draw on examples in the Thresholds Database (Walker and Meyer 2004).

The RSDB includes examples of well-established regime shifts as well as contested and speculative regime shifts. It often takes many years to conclusively establish that a particular change is in fact a regime shift; however, it may be crucial from a management perspective to know that there is evidence suggesting that a regime shift may exist. Furthermore, there are a substantial number of cases where regime shift like phenomena have been observed and described, but have not necessarily been referred to as regime shifts in the literature. The RSDB aims to capture all of these examples that impact ecosystem services. In all cases the level of certainty regarding the existence of a regime shift as well as certainty about the underlying dynamics that cause the shift are clearly recorded. This is based on an assessment of the information and level of agreement in the literature.

The three key criteria for inclusion of both generic regime shifts and case studies in the RSDB are therefore:

1. A large change or reorganization of the SES has been observed or proposed.
2. The change affects the set of ecosystem services provided by the SES, with potential consequences for human well-being.
3. Established or proposed feedback mechanisms exist that create and maintain the different regimes, so that the change is persistent and not readily reversible.

The database has been structured in a hierarchical way so that information can be entered in the form of i) a short summary (case studies only), ii) a more extensive narrative description summarizing the regime shift analysis (both case studies and generic regime shifts), or iii) a detailed regime shift analysis (see next section). The different options enable users to contribute to the database at different levels of detail. The short summary enables additional case studies of generic regime shifts already described in the database to be added easily, as the underlying dynamics and impacts are already captured in the generic description. In most cases the more extensive narrative description is based on a detailed regime shift analysis (see next section), although some users choose to only complete the narrative description. In all cases the information entered in the RSDB draws on material published in the literature.

To ensure data quality, each detailed regime shift is reviewed by either a regime shift researcher or a domain expert prior to publication on the web. We have also included a web form for comment, so that users can provide feedback and updates on the regime shift descriptions, and be engaged in improving the database. To facilitate use of the information in the database and to acknowledge the effort put into the regime shift descriptions, each published entry has a citable reference.

### The Regime Shift Analysis Framework

We have developed a systems-based framework for synthesizing the information in the literature about each regime shift. This framework uses a variety of concepts from systems theory: concepts and approaches from soft systems (Checkland et al. 2006), causal loop diagrams (Sterman 2000), and resilience theory (Bennett et al 2005, Biggs et al. 2015, Scheffer 2009). This framework can also be applied to new contexts where regime shift like phenomena have been observed but not necessarily described as regime shifts.

The systemic portion of the regime shift framework requires the construction of a causal loop diagram (CLD) for each regime shift (generic type or detailed case study) in a consistent fashion. Causal loop diagrams summarize the key drivers and internal feedbacks underlying each regime shift (Meadows 2008), and serves as a visual check on the narrative information in the framework. The level of detail depicted in a CLD always requires choice and judgment, and depends on the purpose of the diagram (Lane 2008).

To enhance consistency across regime shifts we have developed a consistent naming convention for global change drivers (Appendix 1), and rules for the feedbacks and mechanisms that are to be included in a systems diagram. In the case of the RSDB, the CLDs aim to capture the minimum set of variables, particularly the key drivers and feedbacks, which generate the regime shift type dynamics. For generic regime shifts such as eutrophication, there may be several different combinations of drivers and feedbacks that can generate the particular regime shift. In these cases, the CLDs aim to summarize all the proposed and established mechanisms leading to the shift, as the different drivers and feedbacks typically interact and influence one another (e.g. Box 2).

##### Box 2: Regime Shift Analysis for Seagrass Transitions

**Figure B2.**
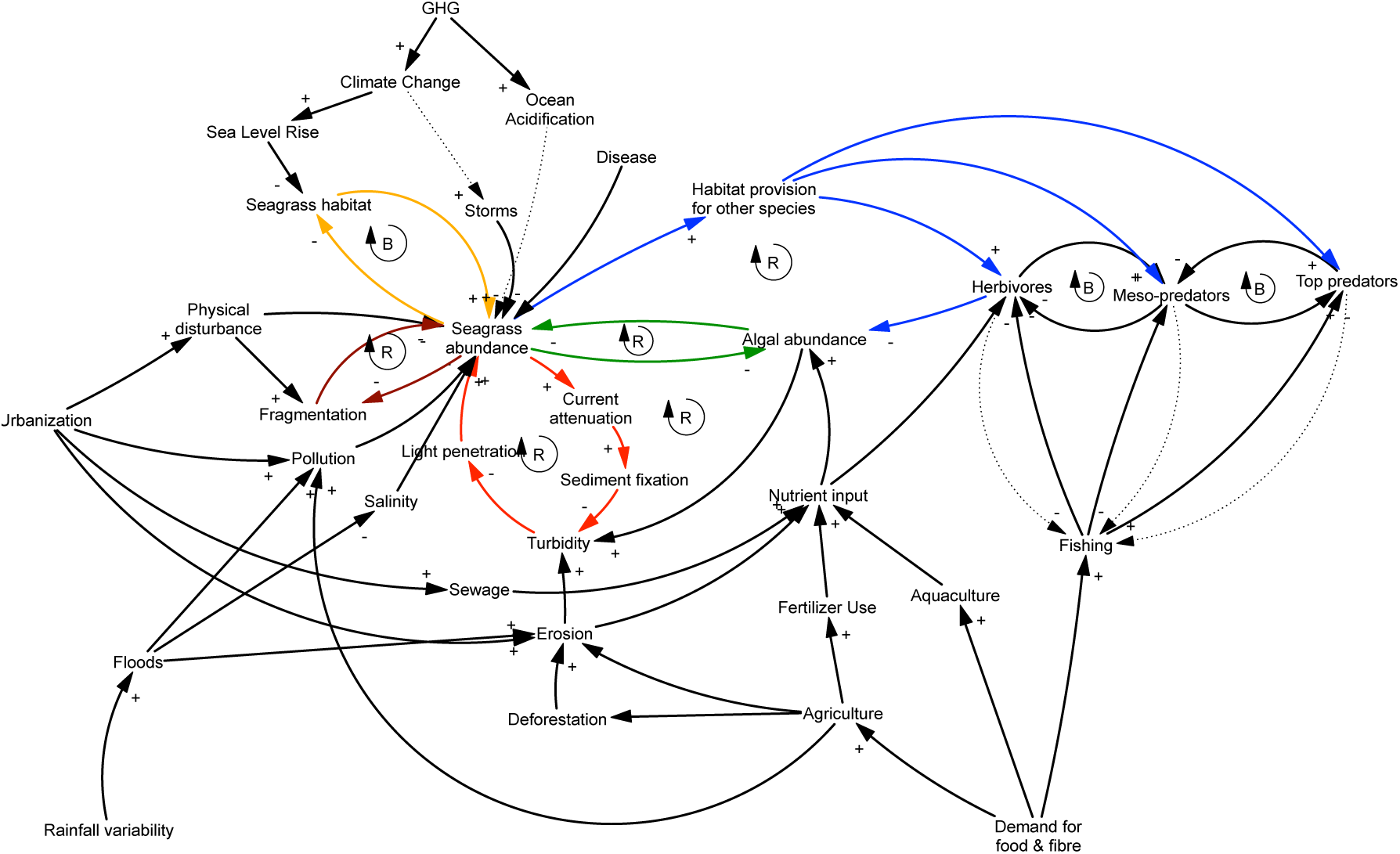
Causal loop diagram. A CLD consist of variables connected by arrows denoting causal influence, with each relationship being either positive (an increase in variable A leads to an increase in variable B or vice versa) or negative (an increase in variable A leads to a decrease in Variable B or vice versa). Closed loops denote feedbacks, which can be either reinforcing (positive) or balancing (negative). Variables that impact on the feedback loops but are not themselves affected by or part of these loops are defined as external drivers (Lane 2008). In the sea grass transitions different feedbacks are highlighted with coloured arrows while drivers relationships are mapped in black. Dashed lines represent causal connections that are uncertain at the scale at which the regime shift dynamics are described.

###### Categorical summary as per database

Regime shifts in seagrass beds are characterised by a collapse of seagrass beds and a transition into either an algae dominated regime or a barren sediment regime. The key drivers are nutrient loading/eutrophication from e.g. agricultural run-off, and overfishing, which both cause slow changes in the system that eventually lead to a sudden collapse of the seagrass regime; or more abrupt shocks like physical disturbance, both anthropogenic and natural, and disease outbreaks that cause direct seagrass decline. Seagrass ecosystems provide valuable ecosystem services such as fishing grounds and coastal protection, which are lost when a shift occurs. Thus human well being can be affected through food and nutrition, livelihoods and economic activity, security of housing and infrastructure as well as aesthetic and recreational values. Once the system has shifted into a new regime it is difficult or even impossible to restore it to its previous seagrass dominated. Therefore ecosystem management should be focused on enhancing resilience in order to avoid a regime shift, e.g. limit nutrient input, reduce physical disturbance and prevent overfishing. This regime shift is typically reported at the local scale (e.g. catchment or community), occur at the time span from months to years, both existence and the mechanisms of the regime shifts are well-established and evidence account for models, contemporary observations, and experiments.

###### Regime shift analysis summary description

Figure 4 depicts the key feedbacks underlying the regime shift dynamcis. The seagrass-turbidity reinforcing feedback (red) shows how seagrass decrease as turbidity increase due to limitation of light penetration. The seagrass – algae competition (green) is a reinforcing feedback that controls for algae and seagrass as they compete for nutrients and space. The herbivory feedback (blue) shows the balancing effect of herbivores on algae that in turn reinforces the growth of seagrass by reducing competition. The habitat feedback (yellow) shows a balancing feedback that denotes density dependence of the population. Direct and indirect drivers relationships are depicted in black. The most prominent direct drivers found in the literature include storms, diseases, physical disturbance such as dredging, nutrient inputs, fishing and aquaculture; while indirect drivers include coastal development and deforestation, green house gases and consequently climate change, ocean acidification and sea level rise. Although specific thresholds were not identified in the literature, it is thought to be related to the amount of nutrients in the water, light penetration, seagrass density and hervibory, as key controlling variables in the feedback loops. Accordingly, leverage points for management include the limitation of nutrients and other pollutants in coastal areas, adaptive management of fisheries paying particular attention to the herbivores functional group, and limiting potential physical disturbance associated to infrastructure development in coastal areas. On a larger scale dealing with climate change is imperative, however managers at the local level might not have the power to influence the physical dynamics of climate or the social dynamics producing green house gas emisions. Another managerial option is transplantation of seagrass and conservation areas.

The second form of the information the regime shift analysis framework contains is a narrative description that complements the systemic description of the CLD. The narrative description includes the following elements:

- *Definition of the system* – Brief introduction to the example, clearly defining the social-ecological system and its boundaries (e.g., lake and its watershed, including the people living in the landscape). The spatial and temporal boundaries of SES are typically “open” and can be defined in different ways depending on the particular focus of the example.
- *Alternate regimes –* Identification and brief description of the different regimes, focusing on what would be seen in the field (e.g., clear water, rooted plants on lake floor, limited agriculture in the catchment). A regime in this context is seen as a particular “configuration” of an SES, characterized by a particular feedback structure and set of functions (Biggs et al. 2012). This is typically the most difficult step in conceptualizing and analyzing each example. To inform ecosystem assessments or management, it is usually sufficient to identify the two or three major alternate regimes, although some of these may comprise several sub-regimes.
- *Feedbacks that maintain each regime –* A description of the key known or proposed feedback processes that maintain each regime, making the regime persistent and difficult to reverse. Each regime is typically associated with the dominance of a particular set of feedbacks that generate and reinforce a particular structure and function of the SES. Different regimes are differentiated by substantive differences in the relative strength of existing feedbacks, or the appearance of completely new feedbacks (Bennett et al. 2005, Biggs *et al*. 2012). For each feedback, the RSDB captures the scale at which it operates (local, regional or global) as well as the level of uncertainty about the feedback (well-established, contested or speculative).
- *Drivers of the regime shift –* A description of the key drivers that cause the system to shift between regimes. These include shocks (e.g., droughts, floods), direct and indirect external drivers, and slow internal system changes. A direct driver directly influences the internal feedback processes underlying a regime shift (but is not itself influenced by the feedback), while indirect drivers alter one or more direct drivers (Nelson et al. 2006). Since drivers depend on the definition of system boundaries, a driver that is direct in one example may be indirect in another. For each driver, the RSDB captures the scale at which it operates (local, regional or global) as well as the level of uncertainty about the driver (well-established, contested or speculative). If the shift can happen in two or more directions, the drivers of each shift are described.
- *Key thresholds –* The levels of key drivers at which a regime shift is triggered, if available in the literature. These correspond to driver levels at which shifts in the dominant feedback processes take place. However, because most regime shifts result from the interplay of multiple drivers, the level of a particular driver at which a regime shift is triggered will depend on the levels of the other key drivers. Consequently, there is usually a whole range of combinations of levels of different drivers that can trigger a particular regime shift. If the shift can happen in two or more directions, the thresholds in relation to each shift are noted.
- *Impacts on ecosystem services* – A description of the ecosystem processes and services that are lost or gained as a consequence of the regime shift. These include provisioning services such as food or clean water, regulating services such as climate regulation or pollination, and cultural services such as recreation and spiritual values (MA 2005). Impacts on biodiversity or ecosystem functions such as primary production or nutrient cycling may also be described here.
- *Impacts on human well-being –* A description of the consequences that changes in ecosystem services trigged by the regime shift hold for human well-being, where human well-being is seen as encompassing multiple dimensions including nutrition, health, livelihoods, security, social relations and freedom of choice (MA 2005). Specific attention is paid to considering which societal groups benefit or lose from particular regime shifts. For this purpose four archetypal groups are considered: Large-scale commercial resource user (e.g., commercial farmers or farming companies, commercial fisherman or companies), small-scale subsistence resource user (e.g. subsistence farmer or fisherman), urban dwellers, and tourists in rural areas. Any impacts on other groups in a particular case are also noted.
- *Leverage points and management options –* A description of the options for i) preventing an undesired regime shift or ii) restoring or encouraging a shift to a more desirable regime. For each leverage point (key SES variable or driver that can be manipulated through a particular management action or intervention), the scale (local, regional, global) and uncertainty (well-established, contested, speculative) is noted, as well as the way in which it influences key drivers and feedback processes to prevent or encourage a shift. Where applicable, differences in management options available to different societal groups are also noted.
- *Uncertainties and unresolved issues –* Many regime shifts are contested or vary in the extent to which their mechanisms are known. This section identifies gaps in knowledge and scientific debates.
- *Key references* - Key literature which readers may refer to for more in-depth information on the particular shift, including literature cited in the narrative descriptions.

The third form of information the RSDB contains is a set of categorical variables that summarizes the information in the narrative descriptions in a way that is comparable across regime shifts. This coding enables easy comparative analyses and provides a mechanism for searching the database in a structured way. This categorical information summarizes the key direct drivers of the regime shift; the land use and ecosystem type in which the regime shift typically occurs; impacts on key ecosystem processes, biodiversity, ecosystem services (provisioning, regulating and cultural), and human well-being; the typical spatial and time scale over which the regime shift occurs; and the reversibility of the shift. In addition, information is given on the evidence in support of the shift (e.g. observations, models, experiments); and the level of confidence about the existence of the regime shift as well as the underlying mechanism (speculative, contested, or well-established). For each regime shift the RSDB lists other regime shifts to which it is connected. For example, marine eutrophication and fisheries collapse are interrelated regime shifts, as each can act as a driver of the other.

Finally, each regime shift entry is accompanied by open source images (diagrams or photographs) illustrating the different regimes. Appendix 2 contains the full data entry template, including details on the possible values for each of the categorical variables.

## CHARACTERIZATION OF THE REGIME SHIFT DATABASE

The RSDB currently contains 27 generic types of regime shifts (Table 1), 10 detailed case studies, and over 300 basic case studies (containing only a short summary). These examples have been contributed by more than 30 different contributors, and have been used by more than 27 Masters students and 14 collaborators to analyze various cases and cross-cutting patterns. The bulk of the database of the database has been developed by researchers and students at the Stockholm Resilience Centre, and the remainder has been contributed by other regime shift researchers. Below we characterize the current state of the database. We focus on the generic regime shifts, and then provide a brief description of the detailed and basic case studies.

**Table 1.** Summary of the generic regime shifts examples currently in the Regime Shifts Database, organized by system type. These constitute among the best-documented environmental regime shifts in the literature. Regime 1 typically refers to conditions with low anthropogenic impacts, while regime 2 refers to conditions with high anthropogenic impacts.

| *REGIME SHIFT* | *REGIME 1* | *REGIME 2* |
| --- | --- | --- |
| **Aquatic systems** |  |  |
| Freshwater eutrophication | Clear water | Murky water |
| Submerged to floating plants | Submerged plants dominance | Floating plants dominance |
| Coastal marine eutrophication | Low nutrient | High nutrient |
| Hypoxia | Normoxia | Hypoxia, anoxia |
| Fisheries collapse | High abundance of commercial fish | Low abundance of commercial fish |
| Marine food web simplification | Predators dominated | Lower trophic groups dominated |
| Bivalves collapse | High abundance of bivalves | Low abundance of bivalves |
| Coral transitions | Coral dominated reefs | Macro-algae, soft corals, corallimorpharians, sponges, urchin barrens |
| Kelp transitions | Canopy forming algae | Turf forming algae, urchin barrens |
| Sea grass transitions | Sea grass | Algae dominated or barren sediments |
| Mangroves transitions | Mangrove forest | Ponds, terrestrial systems, settlements, salt marshes, rocky coast. |
| **Terrestrial systems** |  |  |
| Soil salinization | Low salinity soils | High salinity soils |
| Forest to savannas | Forest | Savanna |
| Peatland transitions | Low productivity & high C accumulation | High productivity & low C accumulation |
| Bush encroachment | Grass dominated savanna | Shrub dominated savanna |
| Tundra to boreal forest | Tundra | Boreal forest |
| Coniferous to deciduous boreal forest | Coniferous forest | Deciduous forest |
| Sprawling vs compact city | Sprawling cities | Dense cities |
| **Spanning aquatic and terrestrial systems** |  |  |
| River channel position | Old channel course | New channel regime |
| Salt marshes to tidal flats | Salt marshes | Tidal or subtidal flat |
| Common pool resource harvesting | High cooperation and resource levels | Overharvesting |
| Indian summer monsoon | Strong monsoon | Weak monsoon |
| Greenland Ice Sheet collapse | Permanent ice sheet | No permanent ice sheet |
| Arctic sea ice loss | Permanent ice sheet | No permanent ice sheet |
| Thermohaline circulation collapse | Strong thermohaline circulation | Collapse of thermohaline circulation |
| West Antarctica Ice Sheet Collapse | Permanent ice sheet | No permanent ice sheet |

### Generic types of Regime Shifts

Ten of the 27 generic regime shifts currently in the RSDB are well-established, both in respect of the existence of regime shifts as well as the underlying mechanism (Figure 1a). A further two examples (forest to savanna, and mangroves collapse) are well-established in terms of their existence, but their underlying mechanism is contested. In general, where the mechanism is not well-understood, there is lower confidence about the existence of a regime shift. The most common forms of evidence in support of the regime shifts recorded to date are models (27 regime shifts), paleological records (20) and contemporary observations (25), with only 14 regime shifts supported by experiments.

**Figure 1.**
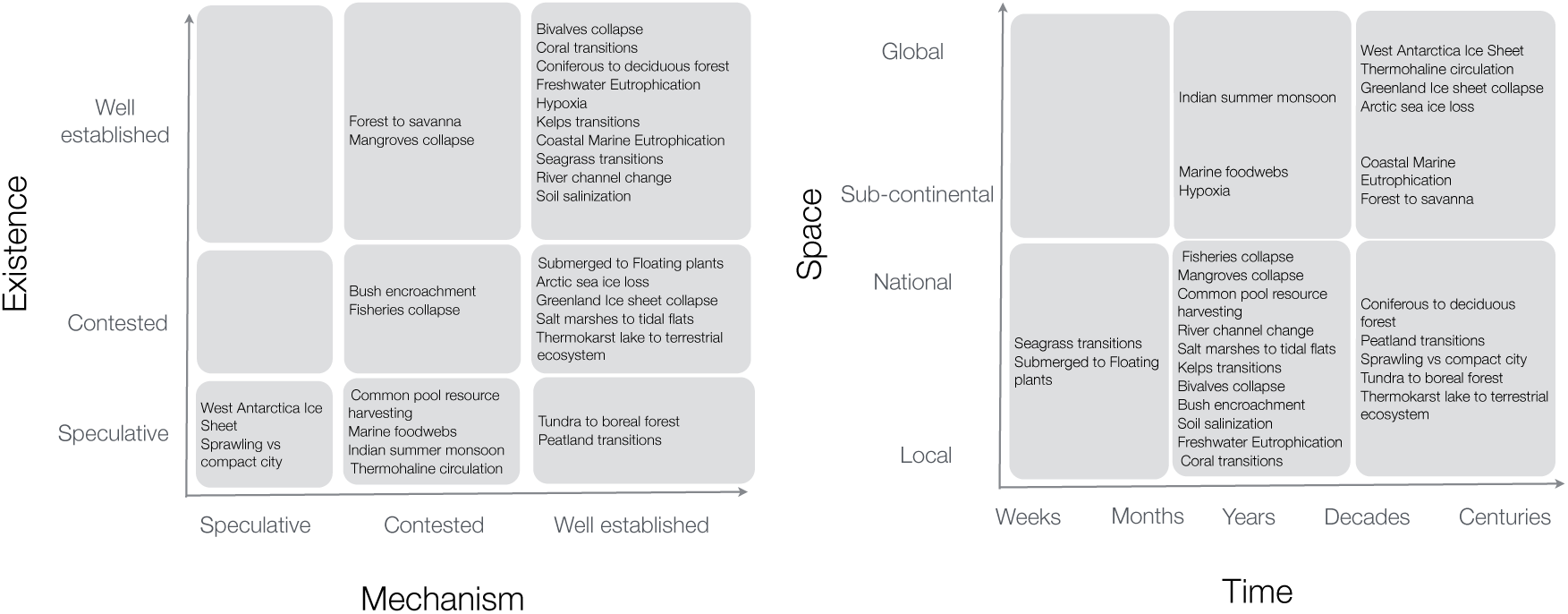
Scale & Evidence. The evidence assessment is shown in a) for both existence of the regime shift and mechanism underlying its dynamics. In b) a Stommel diagram with space vs. time scales for different regime shifts.

More than half of the regime shifts recorded to date occur at the local or landscape scale (20), 13 regime shifts have been reported at sub-continental, 3 confined to national borders and only one at the global (Figure 1b). In terms of timescales, most regime shifts take place over a period of several years to decades (18). These scales are non-exclusive; some regime shifts can occur over several spatial or temporal scales. Twelve of the regime shifts recorded to date are thought to be irreversible on a 100 year timescale, while 17 show evidence of hysteresis.

The regime shifts documented to date have been most commonly found in marine and coastal systems (14 regime shifts), followed by freshwater lakes and rivers (7 regime shifts) (Figure 2a). In terms of land uses under which regime shifts occur, fisheries and large scale commercial crop cultivation dominate (>9 regime shifts each). A large number of regime shifts are also recorded in situations where the land use impacts are primarily off-site, as in the case of marine eutrophication and the transitions from salt marshes to tidal flats. Interestingly, we have recorded relatively few (<6) regime shifts to date under relatively intensive land uses such as urban, small-scale subsistence agriculture, intensive livestock production and mining (Figure 2b).

**Figure 2.**
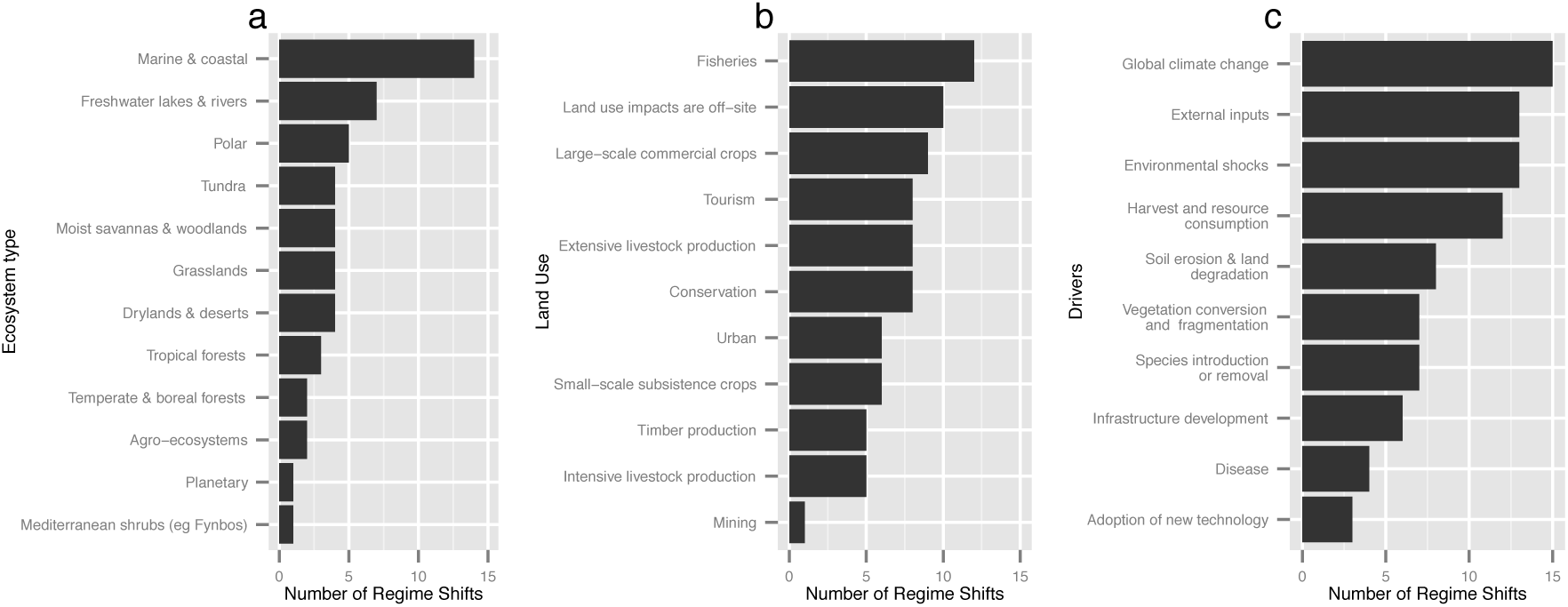
Occurrence and causes of regime shifts. Based on a sample of 27 regime shifts across the globe, a) shows the number of regime shifts that RSDB identifies per ecosystem type, b) summarizes the most common land uses where regime shifts occur, and c) describe the most reported drivers of change.

The regime shift database shows that many different types of drivers cause regime shifts. Global climate change, external inputs (e.g. fertilizers, irrigation), environmental shocks (e.g. fire, floods) and harvest and resource consumption are the most common (>12 regime shifts each). Global climate change is a contributing driver to 15 of the 26 generic regime shifts currently in the database. The least common drivers reported in the examples synthesized to date are adoption of new technologies and disease (Figure 2c).

In terms of impacts, biodiversity is affected by all 27 regime shifts. The most commonly impacted ecosystem processes are primary production (impacted by 17 regime shifts) and nutrient cycling (16 regime shifts) (Figure 3). Climate regulation is the regulating service most commonly affected (15 regime shifts), followed by water purification (12 regime shifts). In terms of provisioning services, fisheries are by far most commonly affected (20 regime shifts), while aesthetic values (21 regime shifts) and recreation (19 regime shifts) are the most affected cultural services. Translating these to impacts on human well-being, livelihoods and economic activity are impacted by 25 of the 27 regime shifts, and food and nutrition by 21 regime shifts. Cultural, aesthetic and recreation values are also commonly affected.

**Fig 3.**
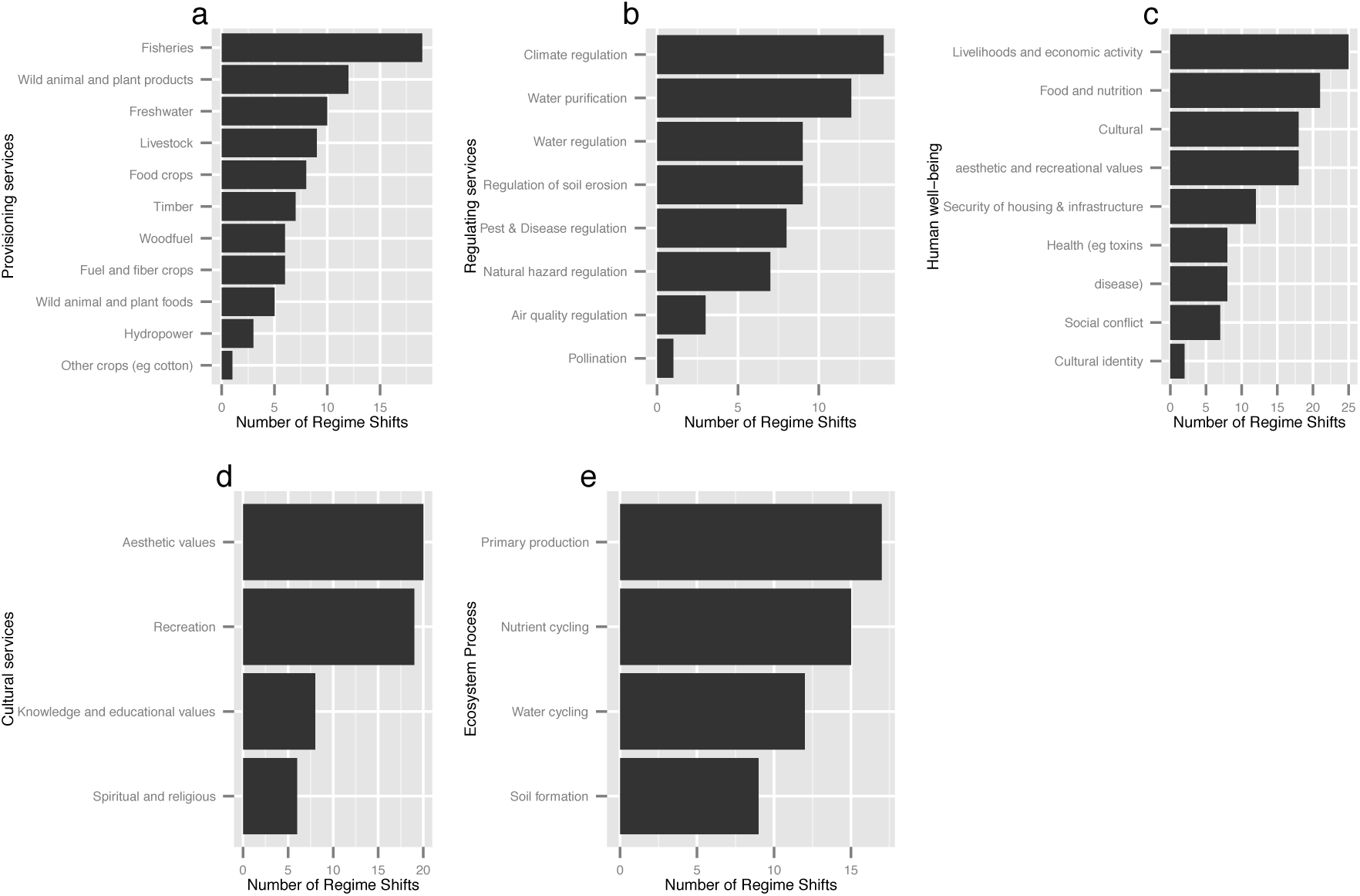
Impacts on ecosystem services and human well-being. Bars describe the number of regime shifts that affect a) provisioning services, b) regulating services, c) human well-being, d)cultural services and e) key ecosystem processes.

### Case Studies

Less effort has been invested to date in compiling detailed case studies of regime shifts. Where these have been undertaken they typically relate to a particular research case that a contributor has been interested to explore in greater depth. As in the case of the generic regime shifts, these examples are also dominated by regime shifts that have been documented in aquatic systems (7 of the 10 detailed cases). The detailed case studies have been a space for conceptual experimentation with regime shifts that have not been framed as such but that are strongly driven by coupled social-ecological dynamics, where the social are not just drivers but also respond adaptively to the dynamics of the ecosystem. Such examples include the Balinese rice production and poverty traps characterizing the Maradi agro-ecosystem. The vast majority of the basic case studies are examples of hypoxia that were compiled from a comprehensive review paper (Diaz and Rosenberg 2008) as part of a student internship.

## DISCUSSION

We have developed a novel, open, empirically based framework for the comparison of regime shifts that combines different perspectives on regime shifts to enable the development of database. This work extends previous work on the analysis of resilience and regime shifts (Holling 1973, Scheffer 2003, Bennett et al 2005). The database builds upon the thresholds database (Walker and Meyer 2004), previous system-based reviews of regime shifts (Gordon *et al*. 2008, Lenton *et al*. 2008, Mård Karlsson *et al*. 2011) to provide the first consistent and extendable database of regime shifts. This database enables researchers to address the need to understand the drivers, prevalence and impact of abrupt change in ecosystems. Below we discuss what we have learned from both the development of the regime shift framework and the regime shift database, before presenting a set of research questions raised by this work.

### Insights from the RSDB

Our database illustrates that the diverse regime shifts that have been documented occur in many different biomes, operate at different scales, and are driven by a wide variety of drivers. This finding suggests that such abrupt changes should be more widely considered in ecological management and assessment, in particular because managing regime shifts requires quite different approaches than managing ecosystems that in which change is consistent and reversible (Carpenter 2003, Scheffer *et al*. 2001), and secondly ecological forecasts that do not consider the possibility of abrupt change are likely to systematically underestimate the impacts of ecological changes. Our database also suggests several surprising patterns in regime shifts: aquatic systems have more regime shifts and regime shifts strongly impact cultural services.

Aquatic systems have more regime shifts than other types of systems. Freshwater, marine and coastal ecosystems have many different types of ecosystems. We don’t know whether this frequency is due to more research in these areas, greater human impact (as most people live near water or the coast), or is more to do with how aquatic environments function. However this difference is interesting and suggests that future research should attempt to better separate these three possible explanations.

We found that regime shifts substantially impact on cultural services, with aesthetic and recreation values being impacted across most regime shifts. Impacts on provisioning and regulating ecosystem services were less widely shared across regime shifts. While regime shifts have frequently been identified as impact provisioning and regulating ecosystem services, such a productivity or water quality, our finding regarding the impact on cultural services is quite novel, and suggests that regime shift research should include consideration of impacts on cultural services. Specifically we believe that conducting a systematic review of cultural services and regime shifts could identify in more detail how cultural services are connected to regime shifts. Furthermore, it suggests that the management and planning connected to cultural services should consider regime shifts.

### What have we learned from the development of the regime shift framework?

Based on the experience of the students and collaborators that have used our framework, it clarifies and speeds regime shifts comparison, but there are still problems with ensuring regime shifts are consistently described and in dealing with uncertain social feedbacks.

The framework’s separation of the description of alternative regimes from the feedback processes and drivers that maintain them clarified and accelerated regime shift analysis, because a common point of confusion for people analyzing regime shifts is a confusion between internal feedback processes and external drivers. The framework’s separation of the behaviour of feedback dynamics from the description of the alternative regimes clarifies this distinction, and by ensuring that feedbacks and divers in the causal loop diagram match the linear blocks of text the comprehensiveness and quality of the regime shift description are both improved. The categorical variables provide a further check, by ensuring that key variables and perspective are included in all the analysis, increasing consistency within and among regime shifts.

Ensuring that regime shifts are consistently described requires coordination among applications of the framework. A regime shift is usually described by an individual based upon a set of literature. While the framework encourages consistency, further constraints are required. We did this by identifying a standard set of drivers and feedback loops that an author can use to define their regime shift, and a review stage where the editors of the regime shift database can work to ensure that processes are explained in a consistent fashion and levels of detail. For example, ensuring that the definition of permafrost, or the driver climate change are consistent across regime shifts. The two level structure of the framework also limits the degree to which a basic description of a regime shift can be inconsistent, and by requiring that this first level be consistent first, provides a strong step towards consistency at the second level. However, because new regime shifts always contain some novelty maintaining consistency in regime shifts still requires iterative revision of the regime shifts across the database.

Another challenge is consistently defining the system boundaries of regime shifts. This is particularly true for less well understood regime shifts. It took over 40 years of experimentation and scientific debate to establish the scale and feedbacks that best explain lake eutrophication or forest to savanna, two relatively well understood cases (Carpenter 2002). However, slightly different system limits bring different takes on what is a driver (direct or indirect), an internal or external process, and can even redefine feedbacks. Many of the controversial regime shifts are often analyzed using inconsistent system boundaries in different studies, for example viewing fishing as internal or external to the regime shift. Contributors to the database face the challenge of integrating different sources of literature and determining which system definitions better match the regime shift dynamic. This problem of system definition also poses a challenge for the consistent analysis of regime shifts.

To make the definition of regime shift system boundaries more consistent we created a consistent hierarchical set of regime shift drivers (Appendix 1), and have defined driver proximity to a regime shift based on connection to key internal feedbacks. These efforts do not solve the problems of consistency and systemic boundaries, but they increase the consistency of the database and increase the ability of people to describe regime shifts in a consistent fashion, by providing consistent criteria to define a system boundary. However, these criteria do not provide a clear answer to whether a particular feedback should be included or not. That decision has to be based on the analysis of existing literature.

A final challenge for the framework is social-ecological feedbacks. Analyzing social-ecological feedbacks is challenging because they are clearly important, but they are often poorly understood. This is especially the case in non-local feedbacks that involve issues such as trade or management, and where feedbacks can often result in learning or policy change that alter a feedback. Understanding these feedbacks is challenging, because often the actors and interactions change over time, and this is particularly true as global trade and markets penetrate in new ecosystems and new forms of information technology accelerate interactions. We strongly believe that new types of approaches to regime shifts are needed to better incorporate uncertain, novel social feedbacks into regime shift analysis, and that this research area of social-ecological regime shifts needs much more investigation, and probably new conceptual models.

### Regime shift research agenda

The regime shift database suggests a number of areas for research. First it identifies gaps in regime shift research, second it suggests a variety of regime shift comparisons, and third it suggests the need to better understand what is special about social-ecological regime shifts.

The regime shift database shows that regime shifts are unevenly studied. A few regime shifts such as eutrophication in lakes, coral reefs, and hypoxia are well studied. However, many regime shifts, such as sprawling versus compact cities or marine food webs, have only been minimally studied. We suspect that the difference in identified drivers among regime shifts is at least partially based on the degree of effort that has been expended studying each regime shifts, with more drivers identified for better studied regime shifts. Consequently, we believe that regime shifts with substantial impact but fewer drivers are regime shifts that merit more study.

The regime shift database enables many different types of comparative analysis of regime shifts. Comparisons across ecosystem type, such as freshwater ecosystems, of region, such as the Arctic, are possible. For example, we conducted a comparison of marine regime shifts (Rocha et al 2015). Similarly comparative analyses of the structure of regime shifts can be conducted, such as analyses of the patterns of regime shift drivers or dynamics. Furthermore, the database could be used to extrapolate the existence of potential unstudied regime shifts, or predict what types of regime shifts may exist in less studied or novel ecosystems.

The third key area of future research suggested by the regime shift database is the analysis of strongly social-ecological regime shifts. Human action now strongly shape most of Earth’s ecological dynamics and humanity increasingly relies of ecosystem services, consequently better understanding social-ecological interactions becomes a key area for regime shift research. However, social-ecological regime shifts that occur in agricultural and urban systems have been have less analyzed than regime shifts occurring in less human dominated ecosystems. Incorporating human action as both driver of and feedback within regime shifts is a challenging task as noted above. There is little understanding whether particular social feedbacks are strong enough and stable over time as to maintain ecological regimes, or if they rather are a source of noise at the time scales at which ecosystem dynamics occur.

The potential importance of social-ecological regime shifts, and the lack of conceptual or empirical understanding of them suggests that they should be a priority area for future research. When drivers and feedbacks are rapidly shifting and evolving a focus on regimes may not be useful. We expect that there are times when the regime shift concepts of alternative regimes, shifts and stabilizing feedbacks will be useful, and others when it is not a good fit and other concept such as pathways or trajectories may be more useful. However, determining the utility of the regime shift concept requires applying it to multiple social-ecological systems. While the regime shift database includes a number of agro-ecological regime shifts, it could be useful to apply the concept to a wider diversity of social-ecological systems. Key questions that could be answered by the analysis of social-ecological regime shifts are: How social-ecological regime shifts differ from ecological regime shifts? How social feedbacks differ from ecological feedbacks? And can the concept of regime shifts be usefully applied to systems strongly shaped by unstable tele-couplings and tele-connections?

We believe that the regime shift database and the regime shift framework can address these and other questions. The regime shift provides a platform for starting point for further regime shift research and an extensible platform for comparative research. While the framework provides a tool for the analysis of new regime shifts, and a starting point for further conceptualization of new aspects of regime shifts.

## CONCLUSION

The regime shift database aims to synthesize dispersed knowledge on regime shifts. The regime shift database can advance regime shift research by enabling researchers to generalize about regime shifts and identify places and conditions that are likely to experience regime shifts. Furthermore, we hope that the database will provide a useful resource of global and regional ecological related assessments, as well as provide a platform the enables the collaboration of researchers, practitioners, and educators interested in better understanding regime shifts, and enables them to developing new understanding of regime shifts.

